# Integrating genome-wide association mapping of additive and dominance genetic effects to improve genomic prediction accuracy in *Eucalyptus*

**DOI:** 10.1101/841049

**Authors:** Biyue Tan, Pär K. Ingvarsson

## Abstract

Genome-wide association studies (GWAS) is a powerful and widely used approach to decipher the genetic control of complex traits. A major challenge for dissecting quantitative traits in forest trees is statistical power. In this study, we use a population consisting of 1123 samples from two successive generations that have been phenotyped for growth and wood property traits and genotyped using the EuChip60K chip, yielding 37,832 informative SNPs. We use multi-locus GWAS models to assess both additive and dominance effects to identify markers associated with growth and wood property traits in the eucalypt hybrids. Additive and dominance association models identified 78 and 82 significant SNPs across all traits, respectively, which captured between 39 and 86% of the genomic-based heritability. We also used SNPs identified from the GWAS and SNPs using less stringent significance thresholds to evaluate predictive abilities in a genomic selection framework. Genomic selection models based on the top 1% SNPs captured a substantially greater proportion of the genetic variance of traits compared to when all SNPs were used for model training. The prediction ability of estimated breeding values was significantly improved for all traits using either the top 1% SNPs or SNPs identified using a relaxed *p*-value threshold (*p*<10^-3^). This study highlights the added value of also considering dominance effects for identifying genomic regions controlling growth traits in trees. Moreover, integrating GWAS results into genomic selection method provides enhanced power relative to discrete associations for identifying genomic variation potentially useful in tree breeding.

## Introduction

Deciphering the genetic basis of complex phenotypic traits is of fundamental importance for understanding biological processes and may ultimately provide information that can help enhance selection in plant breeding programs. Genome-wide association studies (GWAS) is a powerful way to identify putative causal genes or genomic segments that underlie phenotypic variation in plants, particularly for traits with complex genetic architectures (Ingvarsson and Street, 2011; Kruglyak, 2008). Dissection of complex traits have been undertaken in forest genetics to understand the genetic basis of adaptive phenotypes (Ingvarsson et al., 2008; Olson et al., 2013; Wang et al., 2018) or physiological or morphological traits, such as growth or wood properties. For example, Porth *et. al.* (Porth et al., 2013) and later Chhetri *et. al.* (Chhetri et al., 2019a) performed GWAS for wood traits, biomass, eco-physiological and phenology traits in *Populus trichocarpa* with genotyping based on 6.78 million single nucleotide polymorphisms (SNPs). Similarly, a study of *Salix viminalis* identified 29 SNPs that were associated with bud burst, leaf senescence, number of shoots or shoot diameter (Hallingback et al., 2016). In *Eucalyptus,* the earliest GWAS identified 16 markers that were associated with growth and two markers that were associated with lignin traits (Cappa et al., 2013). Recently, 26 quantitative trait loci (QTLs) were identified for productivity and disease resistance using a regional heritability mapping method that helps increase the genomic heritability to 5-15% from 4-6% when using SNPs individually(Resende et al., 2017a; Resende et al., 2017b).

GWAS studies can also provide tools for accelerating the long breeding cycles in tree breeding (reviewed in (Neale and Kremer, 2011)). For example, although many species of *Eucalyptus* display unusually fast growth, breeding cycles aimed at developing elite commercial genotypes still take between 12 to 16 years to complete, since identification of elite genotypes require progeny trials followed by two or more sequential clonal trials (Rezende et al., 2014). However, genomic selection based on genome-wide molecular makers is expected to reduce the time required for completing a cycle of developing elite clones to only 9 years mainly due to the shorter time needed for progeny tests when phenotypes can be predicted from the genomic selection models (Grattapaglia, 2017).

The rapid development in genomics has opened up opportunities to identify molecular markers that are associated with traits of interest and use these marker-trait associations to complement and extend traditional breeding programs. Despite the efforts to discover polymorphisms associated with economically relevant traits, much of the genetic contribution to complex traits in forest trees remains unexplained. One of the main reasons is that GWAS methods normally conduct tests on one marker at a time, for instance using a generalized linear model (GLM) or a mixed linear model (MLM). When dealing with complex traits such as growth and wood qualities, where the effect size of individual loci is likely small to moderate, these methods suffer from limited statistical power to detect loci of small effects (Muller et al., 2017). One potential approach to increase the power and to accurately identify more causal variants is so called ‘multi-locus mixed models’ (MLMM), which simultaneously test multiple markers by including them as covariates in a stepwise MLM to partially remove confounding between tested markers and kinship (Segura et al., 2012). One such method is the ‘fixed and random model circulating probability unification’ (FarmCPU) that performs marker tests using other associated markers as covariates in a fixed effect model (Liu et al., 2016). Optimization across the associated covariate markers using a random effect model is then performed separately. This approach has been reported to simultaneously reduce computational complexity, remove confounding between population structure, kinship and quantitative trait loci, prevent model over-fitting and control the number of false positives (Liu et al., 2016).

Most GWAS analyses to date have been undertaken by implicitly assuming a genetic architecture consisting of additive effects. However, non-additive effects, including dominance (Bruce, 1910), over-dominance (Crow, 1948) and epistasis (Hill, 1982) are known to also play important roles in controlling some traits. One trait where non-additive effects are likely to be pronounced is heterosis, or hybrid vigor, which is the near universally observed phenomenon of phenotypic superiority of hybrid progeny relative to their parents (Charlesworth and Willis, 2009). Not surprisingly, heterosis has been and continues to be of great importance in most plant breeding schemes (Duvick, 2001). To date, a limited number of studies have utilized GWAS methods to dissect the genetic basis of heterotic traits in *Arabidopsis thaliana* and rice. In the model plant *A.thaliana*, dominance and over-dominance of flowering time is a well-studied trait and significant loci from a GWAS were shown to explain as much as 20% of the phenotypic variation in a hybrid population consisting of 435 individuals derived from inter-crossing 30 parents (Seymour et al., 2016). In rice, genome-wide dissection uncovered multiple non-additive effect loci for yield increase (Li et al., 2016; Zhen et al., 2017). For instance, a major QTL, rice heterosis 8 (*RH8*) was found to regulate grain-yield component traits (Li et al., 2016). In *Eucalyptus* hybrids dominance appears to be an important and widespread contributor to many growth-related traits (Bison et al., 2006; Bouvet and Vigneron, 1995; Volker et al., 2008) and ratios of dominance to additive variances exceeding 1.2 have been estimated for growth in *E. grandis* x *E. urophylla* hybrids (Bouvet et al., 2009; Makouanzi et al., 2014; Tan et al., 2017). Such results suggest that there should be ample opportunities to identify SNPs accounting for dominance and/or over-dominance effects in *Eucalyptus* hybrids.

Another genomic-based approach that has become widely used in plant and animal breeding in recent years is genomic selection (GS) or alternatively known as genomic prediction. Unlike GWAS, GS refers to marker-based selection where total genetic variance is captured using genome-wide markers without a prior step of identifying trait-associated markers. GS aims to predict the genetic potential (e.g. genome-estimated breeding values) of breeding individuals without locating genes or QTLs important for the trait(s) of interest. One of most important questions for GS is how to improve the prediction accuracy and methods for accuracy has long been a central research aim in genomic selection. Thus far progress on increasing prediction accuracies have been achieved through the development of new statistical models, more efficient design of training populations, improved quality of phenotypic measurements, a greater number of makers used for model building and by also considering non-additive effects (Grattapaglia, 2017). In this paper we assess methods for improving genomic prediction accuracy by integrating results from GWAS studies into GS to predict the genetic potential of breeding targets. It is well known that using only associated SNPs identified from a GWAS is usually not sufficient for explaining a large fraction of the genetic variation in a trait of interest (the so called “missing heritability” problem, (Makowsky et al., 2011)). However, utilizing GWAS information in the form of associated SNPs, in combination with other types of data has the potential to enhance prediction ability in GS studies (Gowda et al., 2015).

In this study, we present results from a GWAS on growth and wood quality traits, in a breeding population comprising two species of *Eucalyptus* and their hybrids. We also integrate the GWAS results in a GS model with the goal of assessing whether this can help increase prediction accuracies for the traits in question. Specifically, our study has two objectives: first, we implement a state of the art GWAS method that consider both additive and dominance effects for dissecting the genetic architecture of growth and wood quality traits. We also evaluate the proportion of phenotypic variation that explained by significant loci for these two genetic effects. Second, we evaluate how different categories of informative SNPs, selected based on the results from the GWAS, can be implemented in a widely used model for genomic prediction, GBLUP, to estimate variance components and to evaluate prediction accuracies of estimated breeding values.

## Results

### Characters of growth and wood traits

All growth traits were moderately variable at the different assessment ages (Table 1). We observed a lower phenotypic variation for height at 3 years of age, as judged by the coefficient of variation (Table 1). The F1 population underwent selection based on height in order to identify trees to use for genotyping and this selection process likely contributed to the lower phenotypic variation we see in height at 3 years of age. We also observed low phenotypic variation for basic density and pulp yield, which is commonly observed in many wood quality traits. Generally, variation in CBH was greater than in height but both mean and variance for both traits increased as the trees aged. Growth traits generally had low heritabilities (h^2^ < 0.2) whereas wood quality traits showed moderate heritabilities (Table 1). Phenotypic correlations between growth traits were generally positive (0.24∼0.74) whereas basic density was weakly negatively correlated with pulp yield (−0.28). The wood quality traits were generally independent from growth traits (correlations in the range -0.1 - 0.1) (Figure S1). The greatest positive phenotypic correlations were observed between CBH and height assessed at the same age (0.63 and 0.74 for 3 and 6 years, respectively).

**Table 1.** Statistical summary of phenotypes

| Trait | Abbr. | No. obs. | Unit | Mean | CV(%)<sup>†</sup> | h<sup>2</sup> |
| --- | --- | --- | --- | --- | --- | --- |
| Circumference at breast height, age 3 years | CBH3 | 1123 | cm | 61.82 | 13.22 | 0.143 |
| Height, age 3 years | Ht3 | 1094 | m | 22.43 | 9.81 | 0.162 |
| Circumference at breast height, age 6 years | CBH6 | 1104 | cm | 83.80 | 18.67 | 0.186 |
| Height, age 6 years | Ht6 | 985 | m | 28.40 | 13.09 | 0.182 |
| Basic density | BD | 1061 | kg/m<sup>3</sup> | 532.78 | 6.83 | 0.381 |
| Pulp yield | PY | 1039 | % | 49.64 | 8.05 | 0.42 |

<sup>†</sup> CV: coefficient of variance.

### Population structure and model optimization

To examine population structure in the breeding population including both parents and their F1 progenies, we conducted both model-based admixture and fastStructure analyses and principle component analysis (PCA) based on a set of independent SNPs. The admixture analysis could not identify an optimal genetic clustering on account of the minimization of the CV error even for K-values up to K=100 (Figure S2). In contrast, fastStructure suggested an optimal genetic clustering of K=1. Due to the inconsistences between the methods, we repeated the population structure analyses using only the parents, given that the F1 individuals were all obtained through crossings between these parents. Admixture analyses based on the parents alone suggested K=6 minimized the CV error (Figure 1A) and K=6 was also the optimal genetic clustering obtained from fastStructure. The parents were assigned to the six subpopulations according to individual ancestry proportions (Figure 1B). We also performed a PCA to summarize genetic variation among parents and the first six components explained 21.53% of the total genetic variation. Notably, the eigenvalues beyond the first six PCs were relatively small (Figure S3), consistent with the minimum K identified in the admixture analyses. Based on first two principle components, parents can be clearly separated into three clusters and two further sub-clusters can be identified within in each major cluster. Progenies are inferred to be derived from crossing parents either with the different major clusters or between them (Figure 1C) and therefore we used the first six PCs in all subsequent analyses to correct for population stratification.

**Figure 1.**
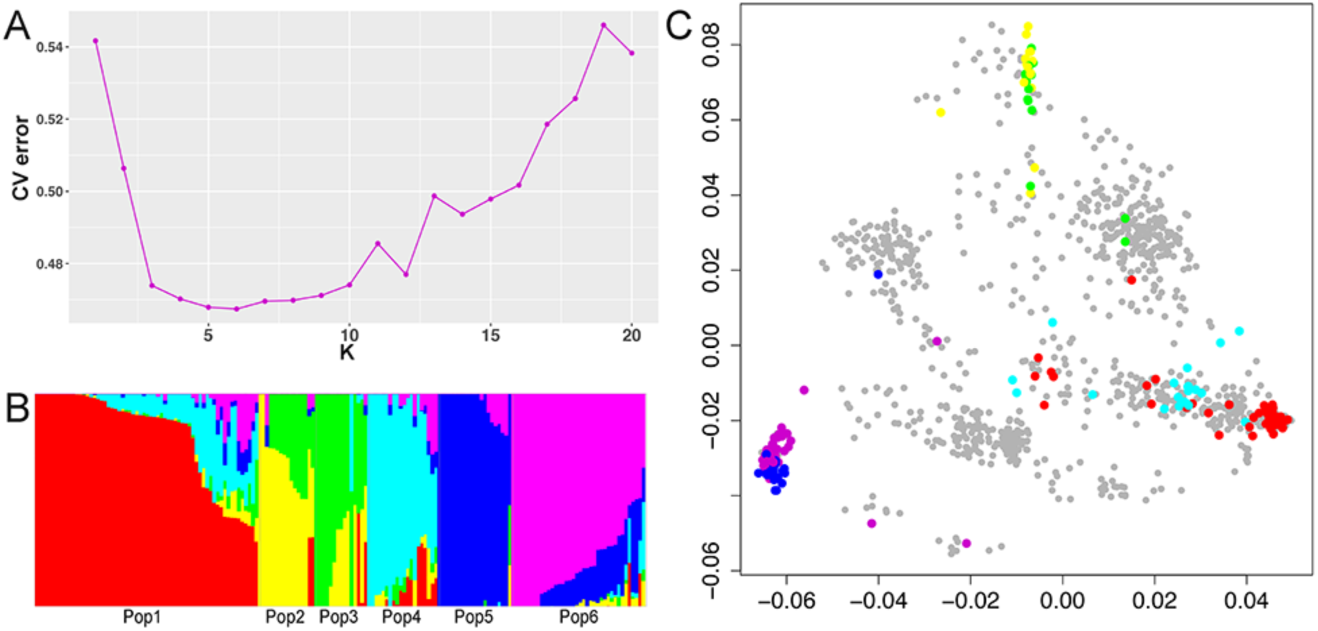
Population structure of parents and F1 progenies. (A) Cross-validation error in the admixture analysis for K varying from 1-20 for the 174 parents. (B) Population structure of parents inferred using admixture for K=6. (C) PCA plot based on genetic covariance among all individuals. Only the first two principle components are shown. The colours used for the parents are in line with the clustering shown in (B), with grey colour denoting all F1 progeny.

### Genome-wide association study for additive effects

We first ran FarmCPU with an additive effect encoding to identify loci with significant additive effects on the different phenotypes. Quantile-quantile (QQ) plots suggest that population structure and kinship relationships were well controlled in the GWAS for the different traits (Figure 2). SNPs with *p*-values < 1.7E-06 threshold were declared statistically significant. Overall, we identified 78 significant SNPs across the six traits and these significant SNPs were distributed across all 11 chromosomes (Figure 2). No significant SNPs were identified for more than one trait, even though both CBH and height show strong genetic correlations across ages. Comparing the number of significant SNPs found for the different traits, growth traits had more significant SNPs than wood traits, with height and CBH at the two different ages having between 14 and 18 significant SNPs whereas we only identified 9 significant SNPs for the two wood quality traits. We generally observe lower phenotypic variances explained by individual SNPs for CBH and height compared to pulp yield and basic density (Table S1). The maximum percentage of phenotypic variance explained by single associated SNP was 2.3% (for pulp yield) and the minimum percentage of phenotypic variance explained by a significantly associated SNP was 0.33% (CBH age 3 years).

**Figure 2.**
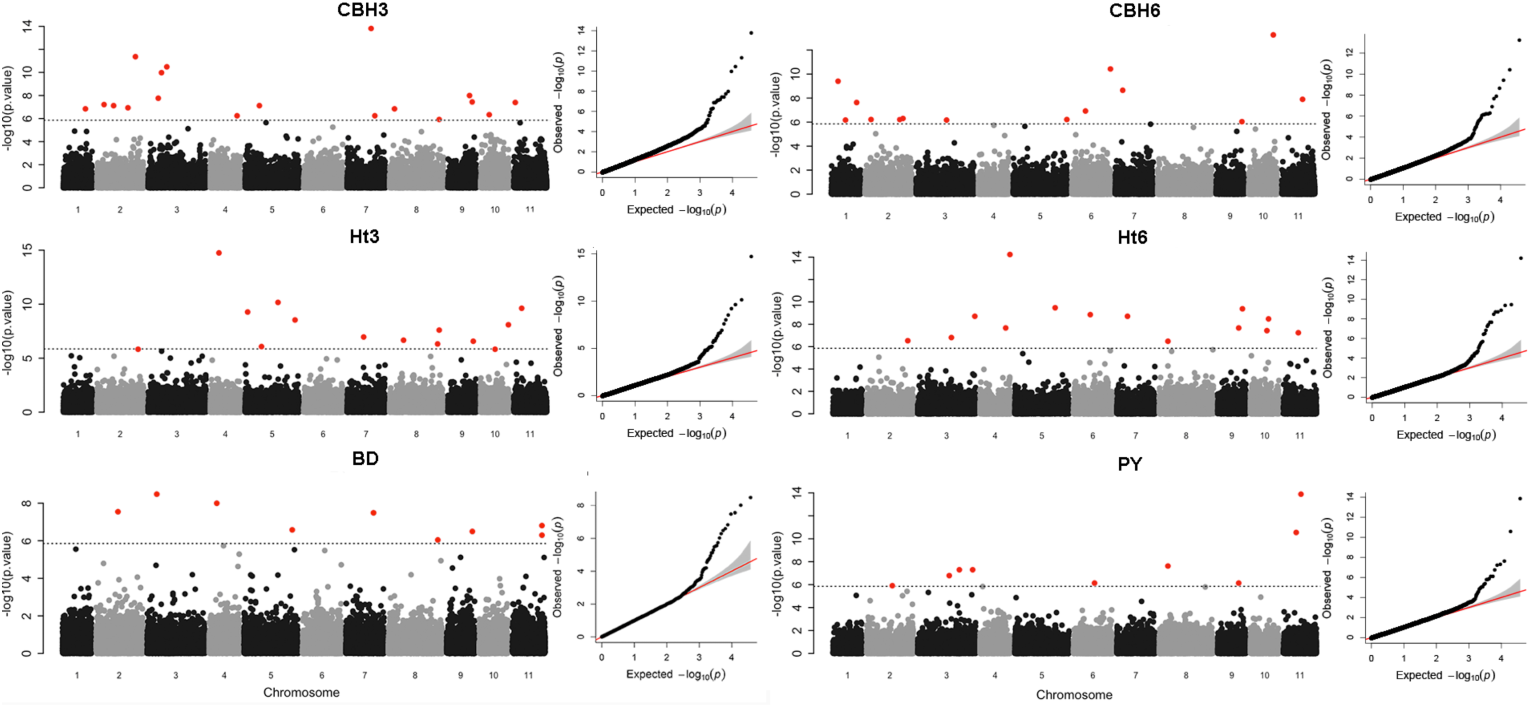
Manhattan plots and quantile-quantile (QQ) plots of the FarmCPU results using an additive effects model. The traits used are CBH and height at age 3 and 6 (Ht3 and Ht6, respectively) as well as basic density (BD) and pulp yield (PY). The Manhattan plots show -log_10_ *p*-values plotted against SNP positions on the 11 *Eucalyptus* chromosomes. Associations reaching genome-wide significance are displayed in red and the horizontal dotted line indicates a Bonferroni-corrected significant threshold of *p* <1.7E-06. The QQ plots for each of the six traits demonstrate the observed versus expected distribution of p-values. The solid red line represents the expected null distribution assuming no associations.

### Genome-wide association study for dominance effects

FarmCPU efficiently controlled the false positive rates due to population structure and sample relationships also when identifying significant loci using dominance encoding (Figure 3, QQ plots). Under a dominance model we detected a total of 82 significant SNPs for the six traits. Height at 3 years old (Ht3) had the greatest number of associations with 19 SNPs displaying significant effects. Fewer associations were observed for the other traits, with between 11 and 15 significant SNPs identified (Figure 3, Manhattan plots). Two significant SNPs, Chr5.40663824 and Chr11.28479550, were found to overlap between CBH and height at age of 6 years. The maximum percentage of phenotypic variance explained by an associated SNP was 4%, a much higher value than found in the additive effect estimations. The smallest percentage of phenotypic variance explained by an associated SNP for the dominance model was of similar magnitude to that observed for additive effects model (Table S2). Comparing significant SNPs identified from the additive and dominance effects models, a total of 10 SNPs overlap between two models for different traits. This result suggest that the two genetic effects are not completely independent. Nine out of ten SNPs that overlap between additive and dominance effects were identified for growth traits and with the remaining SNP observed for pulp yield.

**Figure 3.**
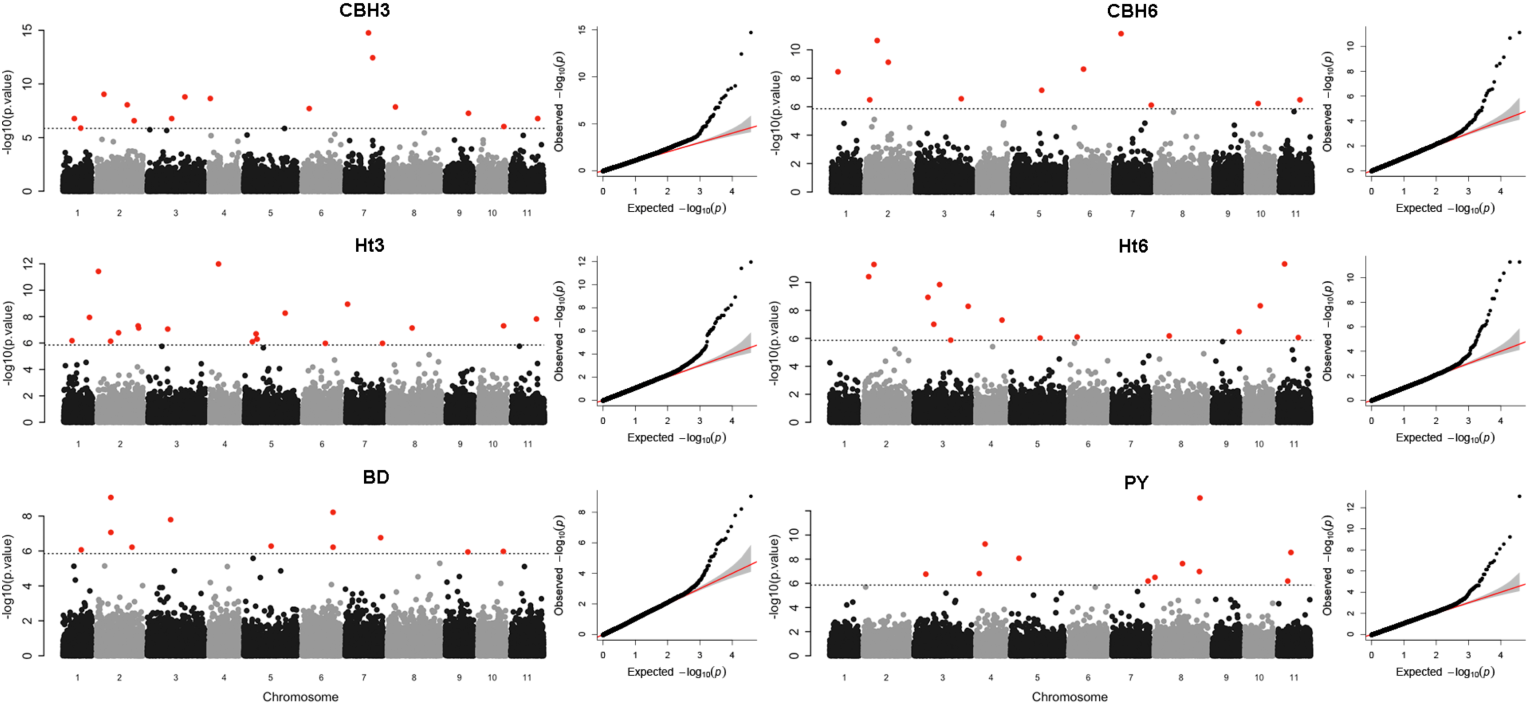
Manhattan plots and quantile-quantile (QQ) plots of the FarmCPU results for the dominance effects model. The traits used are CBH and height at age 3 and 6 (Ht3 and Ht6, respectively) as well as basic density (BD) and pulp yield (PY). The Manhattan plots show -log_10_ *p*-values plotted against SNP positions on the 11 *Eucalyptus* chromosomes. Associations reaching genome-wide significance are displayed in red and the horizontal dotted line indicates a Bonferroni-corrected significant threshold of *p* <1.7E-06. The QQ plots for each of the six traits indicate the observed versus expected distribution of p-values. The solid red line represents the expected null distribution assuming no associations.

### Genome selection by using GWAS results

To confirm the utility of the SNPs identified from the GWAS and to further understand the performance of selecting SNPs for each trait based on the GWAS results, we conducted genomic prediction using four categories of SNPs by using both an additive genetic model (A) and an additive + dominance genetic model (AD). The four categories of SNPs used for the GBLUP analyses were selected from the GWAS results for each trait based on the following criteria: 1) ‘*associated SNPs*’ were identified as significant from the GWAS using the threshold *p*< 1.7E-6; 2) ‘*putative SNPs*’ were identified as significant from the GWAS using a more relaxed significance threshold *p*<1E-3 of each trait; 3) the ‘*top 1% SNPs*’ included the top 1% SNPs for each trait, ranked according to GWAS significance. The rational of this category was to ensure that models utilised the same number of SNPs across all traits. Finally, 4) ‘*all SNPs*’ used all 37,832 available SNPs when building the genomic selection models (Table S3).

The narrow-sense and broad-sense heritabilities were estimated using a modified GBLUP model based on different maker-based relationship matrices calculated using the four SNP categories. As expected, basic density and pulp yield had higher realised heritabilites than growth traits, independent of what category of SNPs that were used for the calculations. Broad-sense heritabilities were higher than narrow-sense heritabilities for most traits, demonstrating that dominance plays an important role in the expression of most traits (Figure 4) and in line with earlier observations in this population (Tan et al., 2018). Comparing heritabilites (h^2^ and H^2^) for the different SNP categories suggest, perhaps surprisingly, that the ‘top 1% SNPs’ category explain more of the genetic variation than any of the other categories, including when all SNPs were used (Table S4). Furthermore, using the ‘top 1% SNP’ set yielded the largest estimates of dominance effects. As expected, using only SNPs that were significantly associated with a trait in the GWAS resulted in lower heritability estimates compared to using all SNPs. Comparing heritability estimates between the ‘putative’ and ‘all’ SNP categories showed that these yielded similar estimates for CBH and height, the ‘putative’ category of SNPs yielding significantly lower heritability estimates than the ‘all’ SNP category for the two wood quality traits (Figure 4).

**Figure 4.**
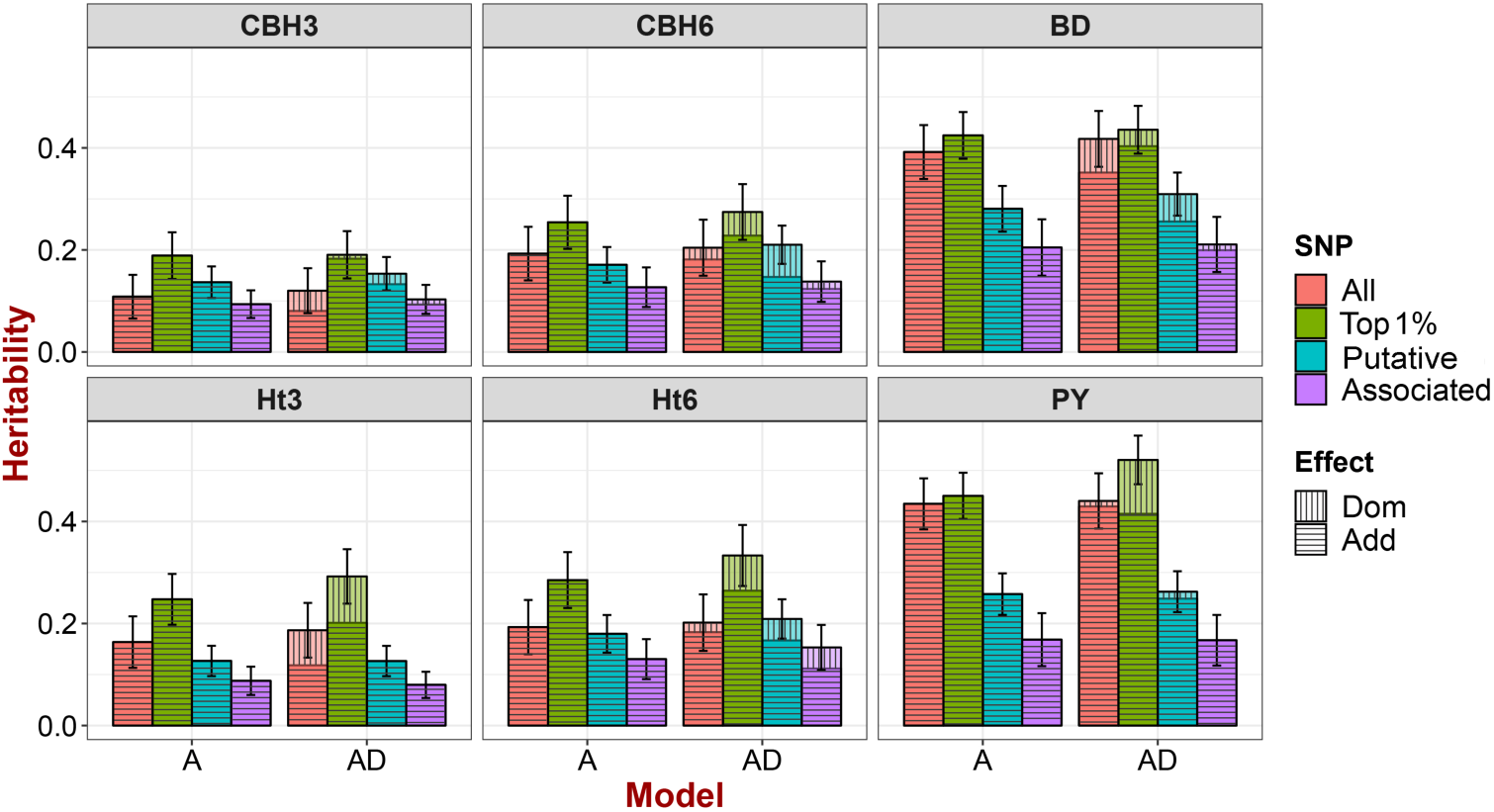
Genomic-based narrow- and broad-sense heritabilities based on an additive (A) or an additive + dominance (AD) model for the four different categories of SNPs used. The coloured bins represent the different categories of SNPs used, with red indicating ‘all’ SNPs (37,832), green indicates the ‘top 1%’ SNPs ranked according to GWAS p-value, cyan denotes ‘putative’ SNPs selected based on GWAS results with *p* < 1E-3 and purple denoted ‘associated’ SNPs selected based on GWAS results using *p* < 1.7E-6. The fill patterns represent different genetic effects. Vertical lines denote additive effects and horizontal lines denote dominance effects. Error bars indicate the standard error of total genetic variance.

**Figure 5.**
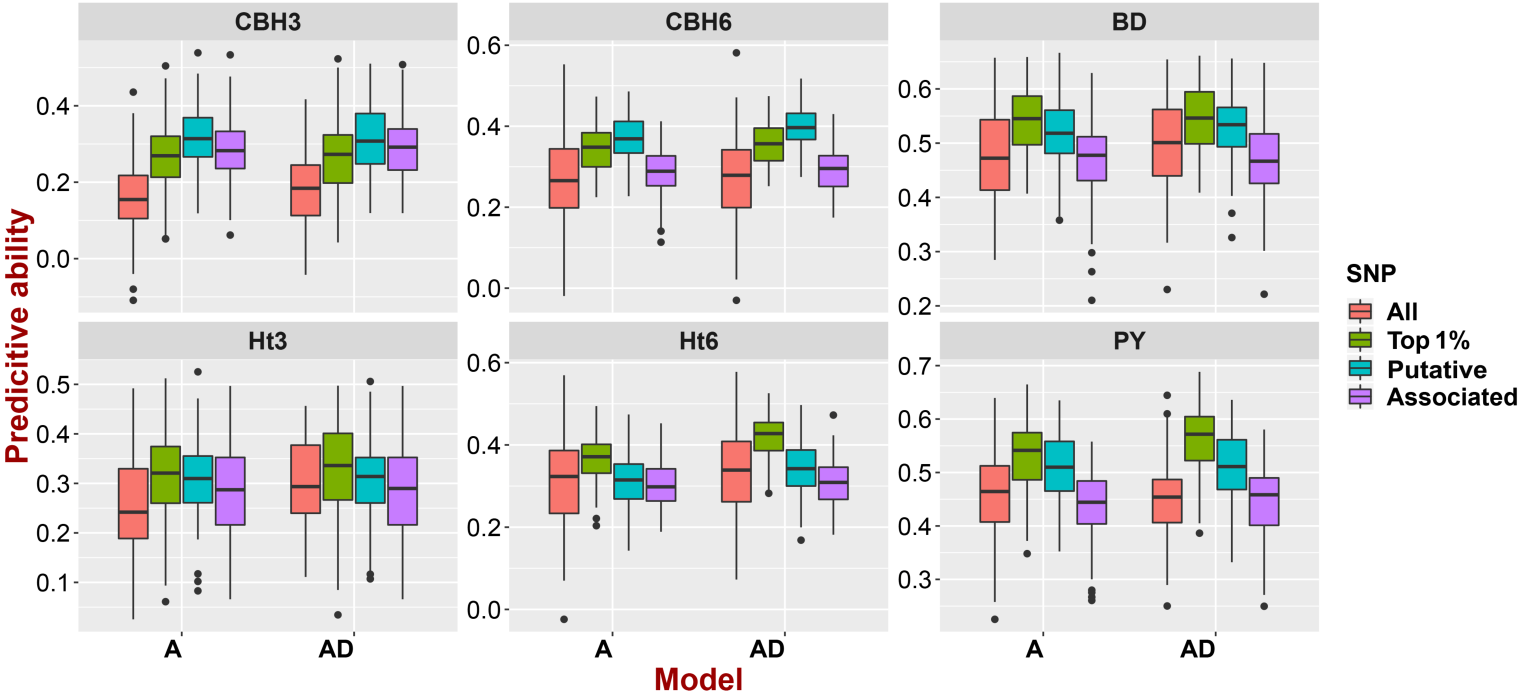
Predictive abilities of the additive (A) and the additive + dominance (AD) genomic prediction models for the four different categories of SNPs. The coloured boxplots display the distribution of predictive ability across 100 replicates of ten-fold cross-validation for the different categories of SNPs. Colours used as in Figure 4.

We further estimated the prediction ability of breeding values for the *A* model and the prediction ability of genetic values for the *AD* model using a ten-fold cross-validation approach. The distribution of predictive abilities for each of the models and SNP, obtained using 100 replications, are displayed in Figure (5). Generally, we observe higher prediction abilities for wood quality traits, in line with the higher heritability values we observe for these traits. The AD model yielded slightly higher prediction abilities than the A model for most of traits. When comparing the different SNP categories, both the ‘top 1%’ and ‘putative’ SNP categories yielded substantial improvements in the predictive ability compared to both the ‘all’ and the ‘significant’ SNP categories. The ‘associated’ SNPs yielded more or less similar results to that obtained using ‘all’ SNPs. Moreover, the ‘top 1%’ SNP category yielded the highest average prediction ability for height, basic density and pulp yield with values being much higher than those calculated from the ‘all’ SNP category (Table S5).

## Discussion

We used GWAS with both additive and dominance effects to dissect the genetic architecture of growth and wood quality traits in hybrid *Eucalyptus*. The method we employed for GWAS, FarmCPU, was able to control the false positive rate induced by both the complex population structure and kinship that characterise our mapping population and efficiently identified significantly associated SNPs for both additive and dominance effects. Using top-ranking SNPs based on the GWAS yielded higher genomic heritability estimates compared to using all available SNPs. We were also able to achieve more accurate genomic prediction results by filtering SNPs based on their associations in the GWAS and this help shed light on future directions in the application of genomic selection in *Eucalyptus* breeding.

### FarmCPU perform superior in GWAS analysis

Most economic traits targeted for breeding in forestry, such as growth and wood properties, are quantitative traits and usually have a complex genetic architecture controlled by many loci of small effect. Here we utilised a recently developed method for the dissection of complex traits, FarmCPU, that has been proposed to efficiently address problems confounding between testing markers and covariates that often arise in GWAS (Liu et al., 2016). Several empirical studies have verified that FarmCPU offers enhanced power for GWAS of complex traits (Vanous et al., 2019; Ward et al., 2019; Zhu et al., 2018). In this study, we identified 78 and 82 significant associations having additive and dominant effects, respectively, with average of 13 SNPs identified per trait studied. These results are more efficient compared to another commonly used GWAS method, Genome-wide Efficient Mixed Model Association (GEMMA). In preliminary analyses we found that GEMMA was also able to control the false positive rate very well, but this came at the price of a relatively low statistical power. GEMMA consequently only identified two significant SNPs for additive effects (Figure S4) and a total of 13 significant SNPs for the dominance effects across traits (Figure S5). The FarmCPU methods therefore appears to be an attractive method that strike a good balance between the identification of false positives and false negatives and thus have good power for the dissection complex traits.

### Dominance effects play important roles in hybrid population

GWAS have traditionally assumed only additive effects of individual SNPs (Bush and Moore, 2012; Marjoram et al., 2014) but here we show the added value of also considering dominance effects for identifying genomic regions controlling growth and wood quality traits. By assessing also dominance effects, we identify an additional 72 associated SNPs across the traits, in addition to the 78 SNPs we identified using additive effects. Furthermore, a considerable proportion of the genetic variation in our hybrid population is attributable to non-additive effects and our results show that the alleles underlying this variation can be identified when dominance effects are explicitly considered in a GWAS setting. Several previous studies have used controlled crosses in crop species, particularly in maize and rice, to identified loci that exhibit dominance effects. For heterosis-related traits, data from maize is frequently cited as supporting the dominance model (Cui et al., 2017; Wallace et al., 2014), while rice has been proposed as a system that supports the over-dominance hypothesis (Li et al., 2016; Zhen et al., 2017). Our approach quantifies the contribution of dominance to the “missing heritability” in a *Eucalyptus* hybrid breeding population and we collectively show that up to 10% of the genomic-based heritability can be explained by associated SNPs that were identified using a dominance model (Table S4).

### The benefit of integrating GWAS results on genomic prediction

Even if we capture a substantially larger number of associated SNPs by considering both additive and dominance effects, a large fraction of the genomic-based hertiabilies (14%-62%) cannot be explained by only considering significantly associated SNPs (Table S4) and these observations are in line with several earlier reports (Chhetri et al., 2019b; Tang et al., 2019; Zhao et al., 2019). Also, when using significantly associated SNPs from the GWAS to investigate the accuracy of genomic predication, we find that this yields no improvement in accuracy, and sometimes even reduced accuracy, compared to predictions based on all available SNPs which mirrors results seen in other similar studies (Gowda et al., 2015; Wallace et al., 2016). Regions identified in a GWAS are consequently not able to explain all of the genetic variation in the traits of interest and this problem is greater for quantitative traits that are controlled by many genes of small effect. These are the traits where current GWAS methods often suffers from insufficient power to detect loci of small effect, unless sample sizes are substantially larger than what is commonly used in most studies of plants.

In order to assess if the GWAS results could be used to enhance genomic prediction in our breeding population, we also tried to identify possible ‘candidate’ SNPs that were not detected as significant using the stringent significant threshold we applied in our GWAS. The rational here is that, as outlined above, most GWAS methods fail to detect loci of small effect but that the GWAS would nevertheless serve as a useful ‘filter’ for ranking SNPs for their possible effects on the traits of interest. We therefore selected two categories of SNPs using two different criteria of relaxed significance and used these to estimate genomic heritabilities and perform genomic prediction. The first category, ‘putative’ SNPs include all SNPs that were found to be associated with the traits of interest based a more relaxed *p*-value (*p*< 1E-3). Using this more relaxed *p*-value we identify between 70 to184 SNPs for the different of trait when combined across the additive and dominance effect models. Using the ‘putative’ SNP category we observed large improvements in the heritability estimates for the growth traits, to the point where almost all of the genetic variation could be explained (Table S4). For wood quality traits, however, about 40% of the genetic variation remain unexplained compared to when using all SNPs for heritability estimation (Table S4). The second category of SNPs we considered consisted of the top 1% of SNPs, ranked by the *p*-value from the GWAS. Using this criterion ensures that the same number of SNPs are used for prediction across the different traits. Surprisingly we were able to explain a substantially greater proportion, up to 174%, of the genetic variation explained when using all SNPs (Table S4). When we performed genomic prediction using these two categories of SNPs we also observe a substantial increase in the prediction ability for all traits compared to predictions based on all available SNPs. This suggests that using all available SNPs introduce noise in the prediction models that negatively affects our prediction ability. Our method for analysing genomic selection and increasing prediction accuracy clearly benefited from integrating results from the GWAS analyses, but the number of associated SNPs that needs to be incorporated depends on the study trait in questions.

### Detection of associations for complex traits in forest trees

Identifying candidate genes underlying growth and wood traits has long been an active area of research in forest trees, such as in *Eucalyptus* (Cappa et al., 2013; Müller et al., 2019; Muller et al., 2017; Resende et al., 2017a), *Populus* (Allwright et al., 2016; Du et al., 2016; Fahrenkrog et al., 2017; Porth et al., 2013) and *Pinus* (Bartholomé et al., 2016; Lu et al., 2017). To ensure good statistical power, both common and rare genetic variants needs to be considered to have a comprehensive understanding of the genetic regulation of complex traits, since many low-frequency variants were identified as associated with growth and wood composition traits (Fahrenkrog et al., 2017). For instance, regional heritability mapping (RHM), has previously been shown to successfully utilise information from both common and rare variants and can therefore capture a larger proportion of the genomic heritability in *Eucalyptus* (Müller et al., 2019; Resende et al., 2017a).

Furthermore, both additive and non-additive effects play important roles in association studies for many traits. Adding dominance effects to a GWAS analysis increase the possibility to identify additional variants that can help capture a greater fraction of the genetic variance (Du et al., 2016; Lu et al., 2017). Other methods, such increasing the sample size using meta-analysis (Müller et al., 2019) or using multi-locus GWAS approaches instead of single marker methods (Fahrenkrog et al., 2017) are methods that also can help increase statistical power in GWAS.

### Putative genes for plant growth and stress response

Among the significantly associated SNPs we observe across additive and dominance effects estimations, we identified a total of 49 candidate genes that have known functions relevant for the traits in question. The details of these genes, including information on the position of associated SNPs and the putative functions of the genes, are summarised Table S6. In general, candidate genes can be separated into two groups, with one group containing genes that have direct functions associated to the morphological formation of different tissues or organs. The other group contain genes related to general responses to abiotic and biotic stress, which, more indirectly, influence plant growth and biomass.

Among the significant SNPs associated with morphology, a number of associations are linked to genes which are related to cell wall biosynthesis. For example, SNP Chr3.46653967 is associated with Ht6 using an additive effects model. This SNP is located on chromosome 3 and encodes a missense variant in a gene coding for a pectin lyase-like superfamily protein (PME). This gene is expressed in stamen and is involved in cell wall loosening and have previously been implicated in floral development (Francis et al., 2006). We also identified a significant SNP on chromosome 11 (Chr11.20479646) which is associated with Ht6 (Table S6). This SNP is located in the gene *Eucgr.K01691* which encodes a homolog to the Arabidopsis alpha-L-arabinofuranosidase 1 (*ARAF*1) gene. Expression of the *ARAF*1 gene is localized to several cell types in the vascular system of roots and stems and the protein is known to be involved in cell type-specific alterations of cell wall structure (Chávez Montes et al., 2008). Many other studies have also identified cell wall biosynthesis related genes from GWAS performed using growth traits in forest trees. Du *et. al.* (2016) identified four significant SNPs that were located in genes involved in secondary cell wall biosynthesis when analysing growth traits in the *Populus* (Du et al., 2016). A SNP associated with volume in *E. pellita* is located in a gene whose function is known to be involved in cell wall cellulose biosynthesiss (Muller et al., 2017). Similarly, Muller *et. al.* (Müller et al., 2019) used a joint-GWAS approach in four *Eucalyptus* breeding populations and identified eight SNPs associated with growth traits that were all linked to genes which were related to cell wall biosynthesis.

Many of the candidate genes putatively related to abiotic and biotic stress show response to adverse conditions. It is perhaps not surprising that these genes show up in our GWAS, as the planting area of the study population alternates between extremely dry (from July to August) and wet (from August to October) conditions in most years, which often leads to stress-induced damage and high incidence of diseases. In line with this, we identified several candidate genes involved in stress response to adverse climate conditions. The SNP Chr2.1760161 is highly associated (*p*-value=3.72E-12) with height at 3 years age in the dominance model. This SNP is located upstream of the gene *Eucgr.B00092*, which encodes a putative *HVA*22 homologue E (*HVA*22*E*). *HVA*22*E* is upregulated to varying degrees in response to cold and salt stress, ABA treatment or dehydration (Chen et al., 2002; Shen et al., 2001). Another variant (Chr3.41941452), associated with CBH at age 6, is located in the upstream region of *High-affinity K+ transporter* 1 (HKT1) gene. *HKT*1 is expressed in root stelar cells and leaf cells (Hamamoto et al., 2015) and provides a key mechanism for protecting leaves from Na+ over-accumulation and salt stress (Berthomieu et al., 2003; Maser et al., 2002). The SNP Chr6.23066996 is associated with CBH at 6 years of age in the additive effects model, is located on chromosome 6 inside the gene *Eucgr.F01775* that encodes catalase 2 (*CAT*2). *CAT*2 controls levels and sensitivity to H_2_O_2_ (Bueso et al., 2007), photo-oxidative stress (Konert et al., 2015) and auxin levels (Gao et al., 2014).

Four of the candidate genes we identify in our GWAS have functions in both morphological formation and stress response. One common SNP (Chr11. 28479550), associated with CBH6 in both the additive and dominance models as well as with Ht6 for the dominance model is located in the vicinity of the gene *Eucgr.*K02133 which encodes a nucleotide-diphospho-sugar transferase (*QUA*1). This enzyme is expressed in vascular tissues and affects homogalacturonan, pectin and hemicellulose cell wall synthesis (Orfila et al., 2005). Recent studies have shown that *QUA*1 also functions in chloroplast-dependent calcium signalling under salt and drought stresses (Zheng et al., 2016). Finally, the SNP Chr4.11644680 is associated with Ht3 and is a synonymous variant located in the *SFR*6/*MED*16 gene which plays important roles in cold- and drought-inducible gene expression (Knight et al., 2009, defence gene expression {Wathugala, 2012 #739) as well as modulating iron uptake (Zhang et al., 2014) in response to cell wall defects (Sorek et al., 2015). These findings suggest that stress resistance also plays an important role in affecting tree growth traits.

## Conclusions

In this study, we have used a GWAS approach in a *Eucalyptus* hybrid population to dissect the genetic basis of growth and wood quality traits by accounting for both additive and dominance genetic effects. Altogether we identify 78 and 82 significant SNPs using additive and dominance models, respectively, with 10 SNPs showing an overlap between the two effect models, suggesting that additive and dominance effects are not completely independent. The associated genes could be grouped into two broad functional categories relating to how they influence tree growth and biomass. One group contain genes associated with morphological formation, such as cell wall biosynthesis, and the other group contain genes related to abiotic and biotic stress responses, such as oxidative, hormone-based and disease-induced stress. These results provide novel targets for possible transgenic or genome editing approaches in the future to directly improve growth and biomass related traits.

We also applied our results from the GWAS in a genomic selection analysis by using different categories of SNPs selected based on the GWAS results and used them to evaluate genomic-based heritabilities and predictive abilities. Our results show that prediction abilities of the estimated breeding values improved for all traits when using SNPs selected based on the GWAS results. Integrating GWAS results into genomic selection thus appear to be a promising avenue to increase the efficiency of genomic selection in forest breeding.

## Experimental procedures

### Populations, phenotypic and genotypic data

A total of 1123 *Eucalyptus* individuals were used in this study, including 90 *E.grandis*, 84 *E.urophylla* parents and 949 F1 progenies derived from a random mating design that has previously been described (Tan et al., 2017). Briefly, F1 individuals were identified to be comprised of inter- and intra-crossing of the two parental species. Of the 949 F1 individuals, 57% were interspecific *E.grandis* × *E.urophylla* hybrids, 21% were intraspecific *E.grandis* × *E.grandis* progeny and 22% were intraspecific *E.urophylla* × *E.urophylla* progeny (Tan et al., 2018).

The phenotypic and genotypic data utilized in this study has been previously described in detail (Tan et al., 2017). The phenotypes include height and circumference at breast height (CBH), where F1 individuals were evaluated at ages three and six and the pure species parents were evaluated at age five. In addition, we obtained data on two wood quality traits, basic density and pulp yield, that were assessed at age five. Genotyping was performed using an Illumina Infinium EuCHIP60K SNP chip that contains probes for 60,904 unique SNPs (Silva-Junior et al., 2015). Across the 1123 individuals, 37,832 SNPs were retained after quality-control based on call rates (>0.7) for both SNPs and samples and following filtering based on minor allele frequencies (>0.01) and deviations from Hardy-Weinberg equilibrium (>1e-7). Any missing data remaining in the 37,832 SNPs were subsequently imputed using BEAGLE 4.1 (Browning and Browning, 2007).

### Phenotypic data analyses

Phenotype data for the parental and F1 population were adjusted separately to minimize environmental variation by fitting a mixed linear model for each trait:

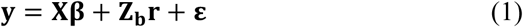

where **y** is the vector of phenotypic observation, **β** is the vector of overall mean as fixed effect, **r** is the vector of random replication effects following 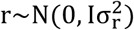, where 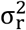 is the replication variance, **ε** is the vector of random residual effects. **X** and **Z**_b_ is design matrix for **β** and **r**, respectively. For the F1 population, the residual variance-covariance matrix is 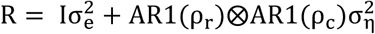, where AR1(ρ_r_) and AR1(ρ_c_) are autoregressive correlation matrices for the row model (autocorrelation parameter ρ_r_) and column model (autocorrelation parameter ρ_c_), respectively. 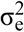 is the independent residual variance while 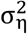 is the spatial variance. For the pure parental species, we fitted the model in Equation 1 by setting the residual matrix 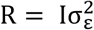 since spatial coordinates and position information were not available for these individuals. All mixed-linear model analyses were performed in ASReml 4 (Gilmour et al., 2015). Phenotypes of individuals from the F1 and parental populations were adjusted for random block effects (r) and spatial effects (s), respectively.

The heritability (h^2^) was estimated using a mixed model **y** = **Xβ** + **Zu** + **ε**, where **y** represents the adjusted phenotypes of single trait, **β** is the vector of fixed effects, including overall mean and age difference. **u** is a vector of random additive or dominance genetic effect of individuals with a normal distribution, 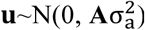, **A** being the revised, pedigree-based genetic relationships among individuals (Tan et al., 2018); and **ε** is a heterogeneous random residual effects represented different experimental sites. **X** and **Z** are incidence matrices for **β** and **u**, respectively. We obtained restricted maximum likelihood (REML) estimates of σ_a_^2^ and **ε**, and estimated 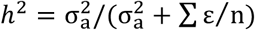.

### Population structure, kinship analysis and GWAS

In the association analyses, confounding effects of population structure and kinship between individuals need to be accounted for. Population structure (Q) was estimated using a model-based clustering and through principle component analysis (PCA) using 13,245 independent SNPs obtained by LD-pruning the original SNP data set and including only SNPs that have pairwise linkage disequilibrium (LD) values (*r*^2^) less than 0.2. Model-based clustering was implemented using the software *admixture* v.1.3.0 which infer population structure by estimating individual admixture proportions using multi-locus SNP data through a maximum-likelihood method (Alexander et al., 2009). The number of ancestral populations (P) was varied from 1 to 100 when using admixture and *fastStructure* (Raj et al., 2014). Five-fold cross-validation (CV) was performed to choose the optimal P value in admixture. A PCA was also performed using the *smartpca* program in *eigensoft* v6.0 to estimate individual ancestry proportions (Price et al., 2006).

GWAS was conducted using a recently developed method, FarmCPU, which explicitly takes into account the confounding that exists between covariates and test marker by using both a fixed effect model and a random effect model (Liu et al., 2016). The results from the PCA and the kinship matrix were used as covariates in FarmCPU to account for population structure and relatedness among samples, respectively. We ran the GWAS using the R package *FarmCPU*. False positive errors due to multiple testing were controlled by an adjusted Bonferroni method, *simpleM* (Gao et al., 2008). This method infers the number of independent SNPs by filtering on LD and performs a standard Bonferroni correction to correct for multiple testing based on the number of ‘independent tests’ performed. For the present data a *p*-value < 1.7E-06 was selected as a cut-off to indicate significant associations.

### GWAS for additive and dominance models

We conducted GWA analyses in FarmCPU using either an additive or dominance encoding of genotypes. For the additive encoded data, the homozygous major allele was encoded with 0, the heterozygous genotype with 1 and the homozygous minor allele with 2. For the dominance encoding, both homozygous minor and major alleles were encoded as 0 whereas the heterozygous genotype was encoded as 1 (Seymour et al., 2016).

### Genomic selection (GS) with different informative SNPs

Genomic selection (GS) models were constructed based on four different categories of SNPs using the Genomic Best Linear Unbiased Prediction (GBLUP) method. Details on how the genomic-based additive and dominance relationship matrices are estimated have been previously described in detail in Tan *et al*. (Tan et al., 2018). Here we focus on the details of how the four categories of SNPs we have used in all subsequent analyses were selected. The four categories of SNPs employed for estimating the additive and dominance relationship matrices were: 1) ‘*associated SNPs*’ which contain only SNPs that were identified as significantly associated with the corresponding trait at the Bonferroni-adjusted *p*-value <1.7E-6; 2) ‘*putative SNPs*’ are all SNPs that were significant in the GWAS for the corresponding trait using a more relaxed *p*-value threshold (*p*< 1E-3) in order to capture also possible causal SNPs that do not reach significance using the more stringent criteria in the original GWAS; 3) The ‘*top 1% SNPs*’ category use the top 378 SNPs for each trait in the GWAS ranked after *p*-value in the GWAS in order to evaluate the same number of SNPs across different traits when building genomic selection models; and finally 4) ‘*all SNPs*’ which use all of the 37,832 SNPs available and is therefore identical to the models originally used in Tan et al (2017) and Tan el al (2018).

Two separate GBLUP models were evaluated that included i) either only additive (*A*) or ii) both additive and dominance (*AD*) genetic effects using the four SNP categories described above to create marker-based relationship matrices. The *A* and *AD* models have been well described earlier in Tan *et. al.*(Tan et al., 2018). The genomic-based narrow- and broad-sense heritability (*h*^2^ and *H*^2^ respectively) were calculated after fitting each model across the different traits. Narrow-sense heritability in the *A* model was estimated as 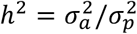 and the broad-sense heritability of *AD* model was estimated as 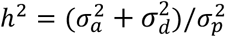, where 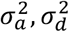 and 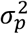 represented the estimated additive, dominance and phenotypic variance, respectively. The prediction ability was estimated for all models and relationship matrices using a ten-fold cross-validation scheme where 100 replications was implemented to evaluate the prediction accuracy of the different models. For each replication, the dataset was randomly divided into 10 subsets and nine out of the ten partitions were used as the training population to fit a model using both phenotypes and genotypes, while the remaining partition was used as the validation set where phenotypic data was removed and then used to predict breeding values or total genetic values for the model in question. The predictive ability of the model was evaluated by estimating the correlation between phenotypes and breeding/genetic values, 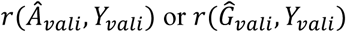.

### Assigning significant SNPs to putative candidate genes

Genes within ±5kb away from a SNP that was significantly associated with a measured phenotypic trait were extracted from the *E.grandis* v2.0 reference genome (BRASUZ1) in Phytozome (www.phytozome.net) using SnpEff v4.2 (Cingolani et al., 2012). The 5kb window threshold used was based on the distance over which LD decays in this population (Tan et al., 2017). The putative functions of these candidate genes were determined based on their homology to functionally characterized genes in *A. thaliana* (TAIR).

## Acknowledgements

The study has partly been funded through grants from Vetenskapsrådet and Kempestiftelserna to PKI. BT gratefully acknowledges financial support from the Umeå Plant Science Centre (UPSC) “The Research School of Forest Genetics, Biotechnology and Breeding”. The research was conducted using resources from the High Performance Computing Centre North, Umeå (HPC2N). The authors are grateful to Yousry El-Kassaby for comment on an earlier version of the manuscript.

