## Supplementary Figures for "Integrating genome-wide association mapping of additive and dominance genetic effects to improve genomic prediction accuracy in *Eucalyptus*"

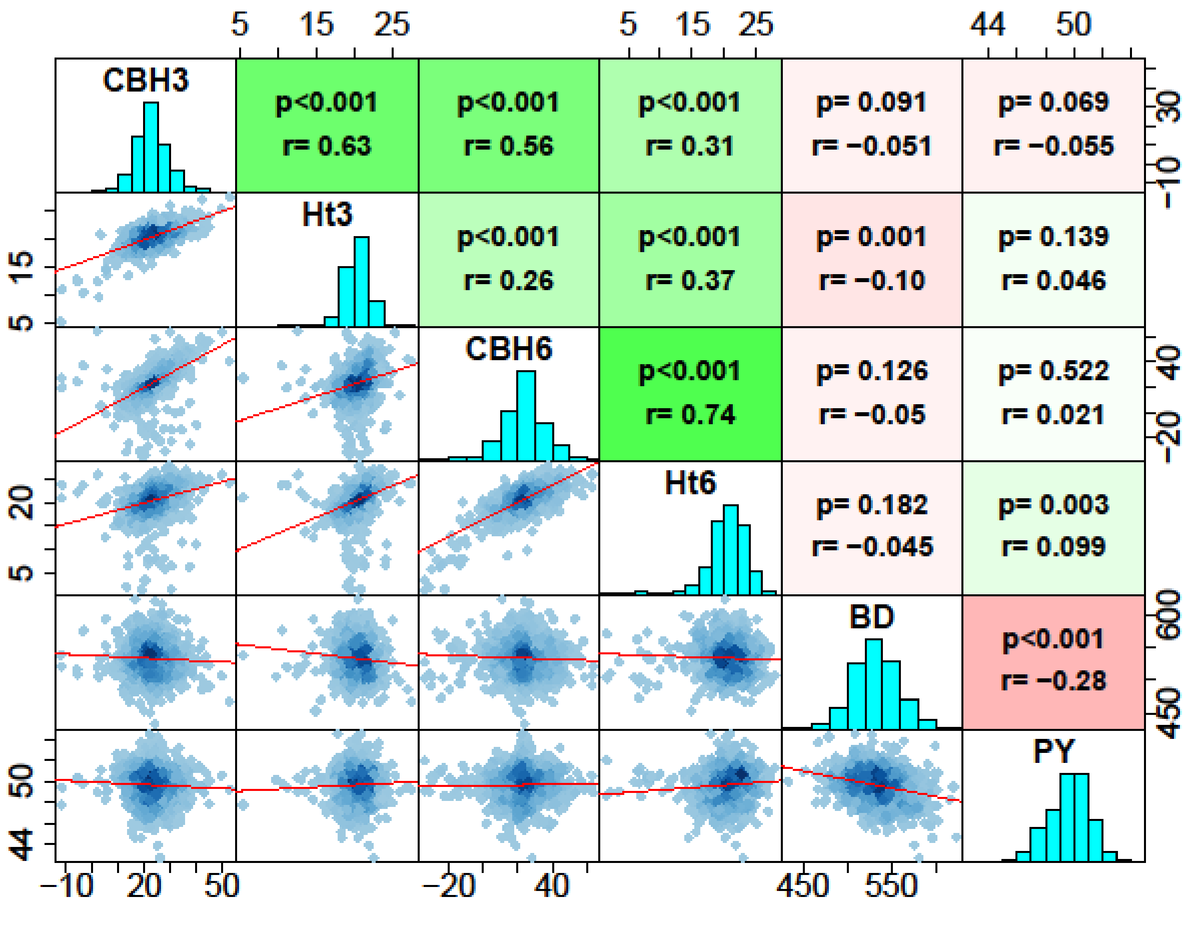


Figure S1. The distribution and correlation of phenotypes. Scatter plot (lower off-diagonal) and correlations with probability values (upper off-diagonal; $H_0$: r=0) for adjusted phenotype among the study traits. Diagonal is the histograms of the distribution of the study phenotypes.

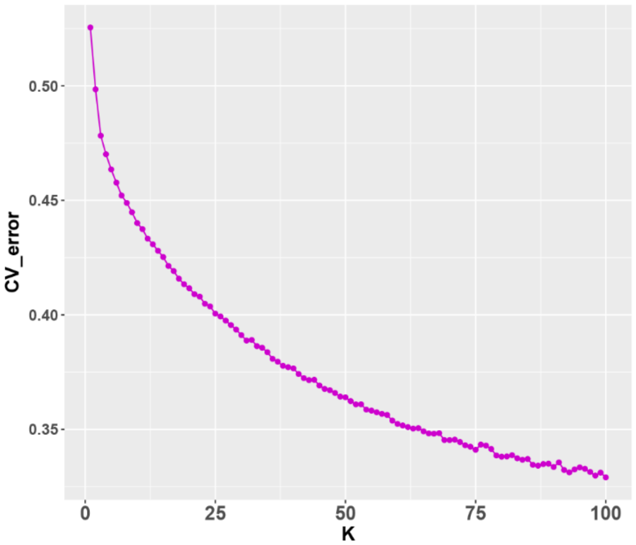


Figure S2. The cross-validation error of admixture analysis. K in x-axis represent the test of clustering the population of parents and F1 progenies into the number of groups. Here we tested K from 1-100. The smaller in CV_error means the number of the cluster is the best.

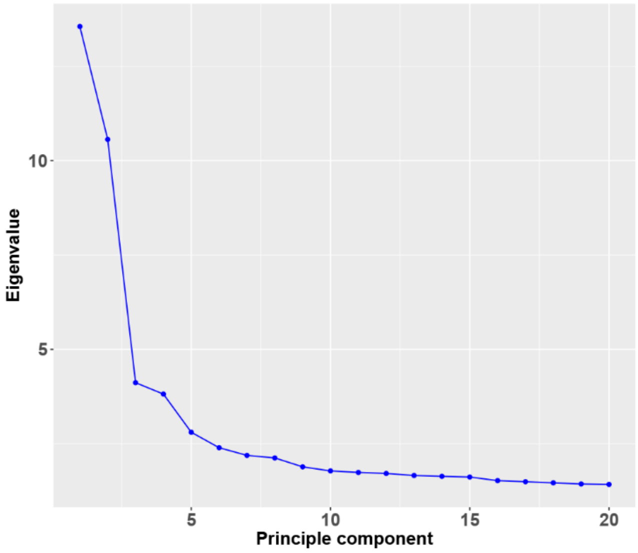


Figure S3. The eigenvalue of the principle component analysis for testing population structure. X-axis represent the number of principle component were used and the Eigenvalue of each component.

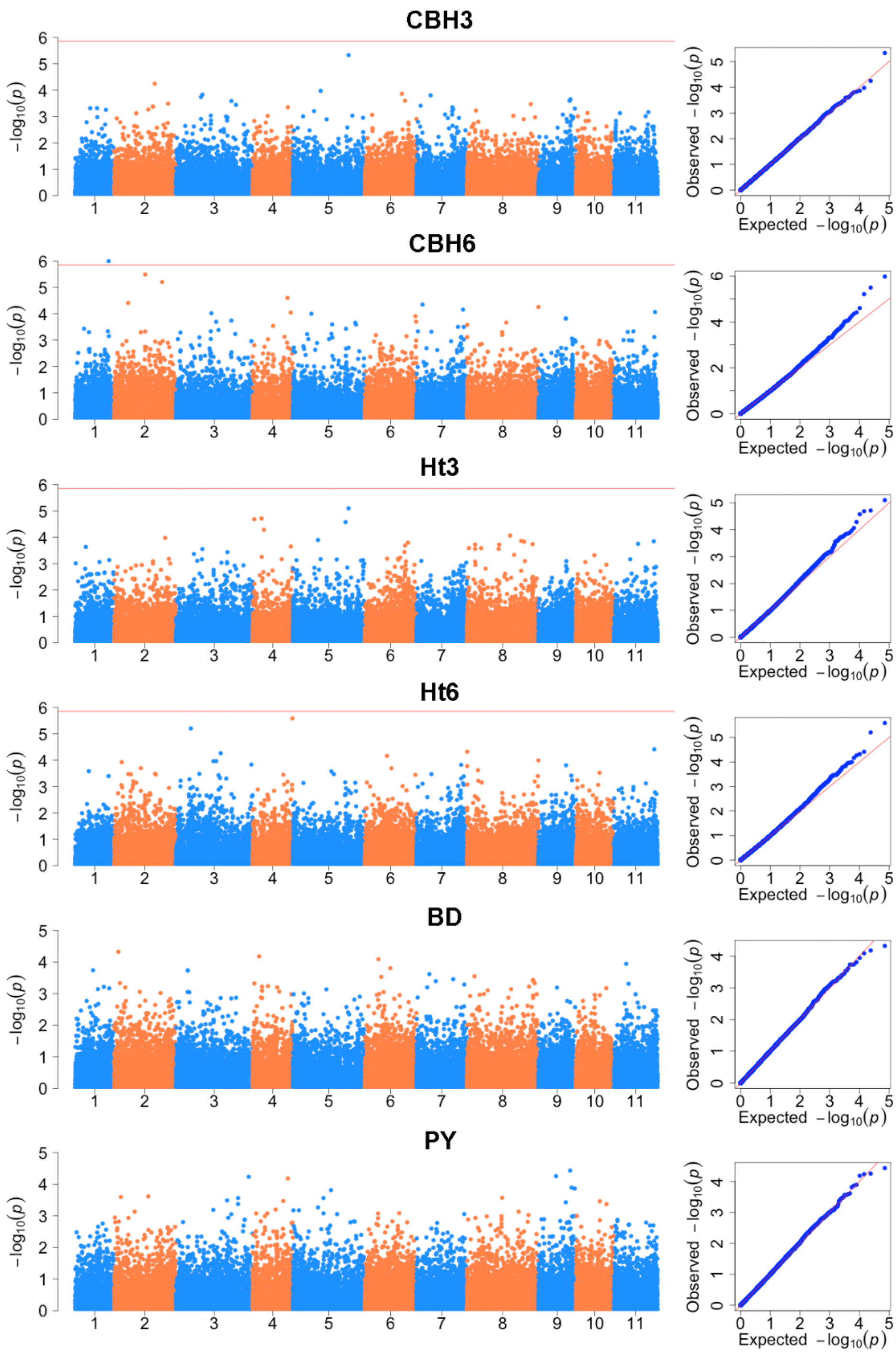


Figure S4. Manhattan plots and Quantile-Quantile (QQ) plots represent the GEMMA results of estimating additive genetic effects. CBH and height (Ht) at 3 years and 6 years, respectively, basic density (BD) and pulp yield (PY) traits are participated the analyses. Manhattan plots indicate negative log10 P-values from a genome-wide scan are plotted against SNP positions of 11 chromosomes. The horizontal red lines indicate the significant threshold of P-value<1.7E-06. QQ plots of each trait showing expected null distribution of P-value vs. distribution of observed P-value, the solid red line represents the null distribution abline assuming no associations.

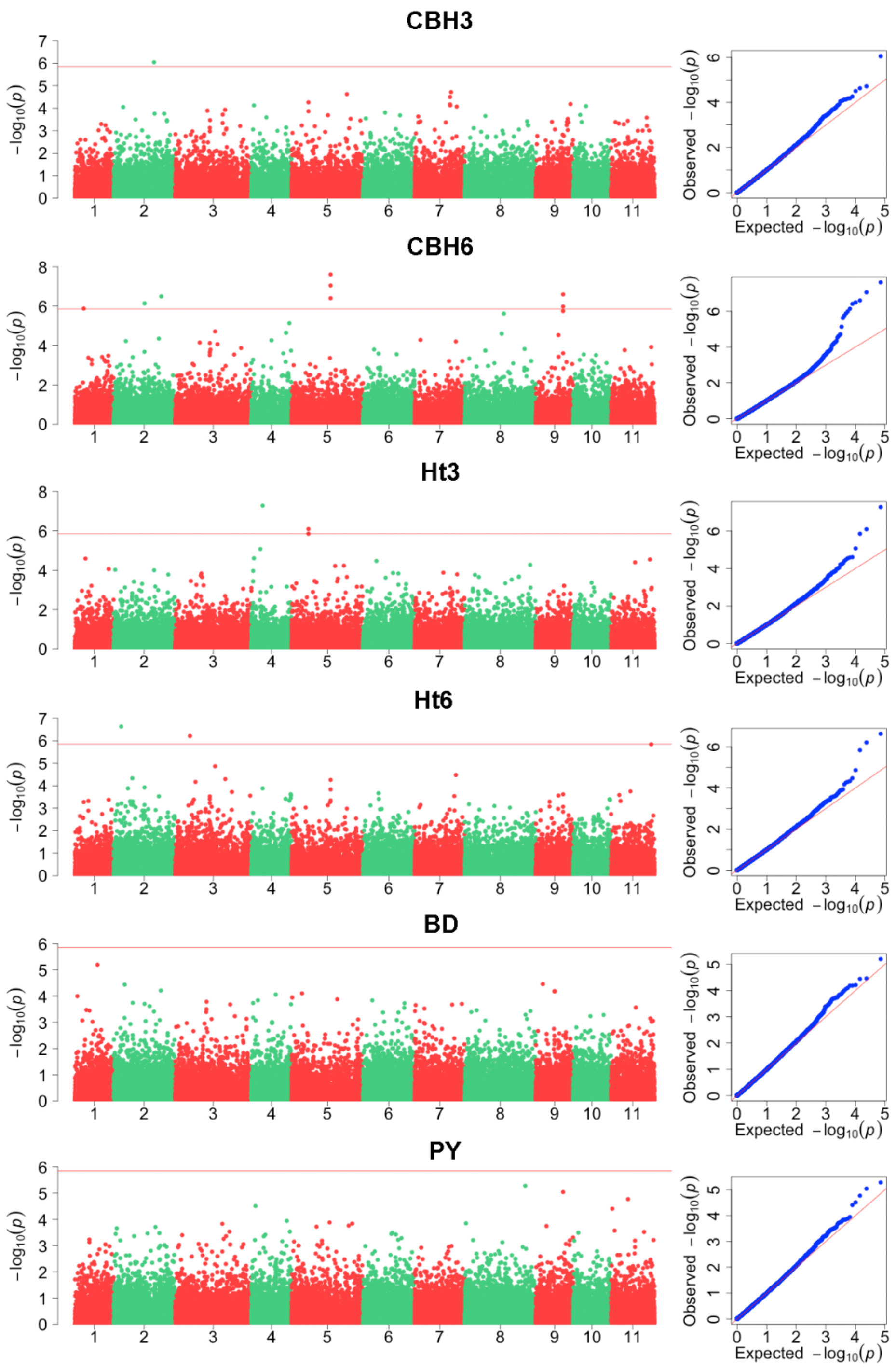


Figure S5. Manhattan plots and Quantile-Quantile (QQ) plots represent the GEMMA results of estimating over-dominance genetic effects. CBH and height (Ht) at 3 years and 6 years, respectively, basic density (BD) and pulp yield (PY) traits are participated the analyses. Manhattan plots indicate negative log10 P-values from a genome-wide scan are plotted against SNP positions of 11 chromosomes. The horizontal red lines indicate the significant threshold of P-value<1.7E-06. QQ plots of each trait showing expected null distribution of P-value vs. distribution of observed P-value, the solid red line represents the null distribution abline assuming no associations.
